# Breaking Bad: Exploring & Complementing the Effects of the DNASE1L3 p.Arg206Cys Variant on Cell-Free DNA from an Isogenic Cell Line Model

**DOI:** 10.1101/2025.08.05.668479

**Authors:** Kavish A.V. Kohabir, Jesper A. Balk, Lars O. Nooi, Dimitra Papaioannou, Rob M.F. Wolthuis, Erik A. Sistermans, Jasper Linthorst

## Abstract

DNASE1L3 is a key endonuclease, essential for proper fragmentation and clearance of cell-free DNA (cfDNA). The p.R206C common variant impairs DNASE1L3 secretion and activity, causing aberrant cfDNA fragmentation and therefore affecting liquid biopsy-based screening and diagnostics. Existing studies on DNASE1L3 relied on resource-intensive murine models or plasmid-based overexpression, which do not accurately represent native expression. To address this, we developed an isogenic HEK293T cell line model by using CRISPR Prime Editing for endogenous expression of *DNASE1L3*^*R206C*^. We analyzed the cfDNA composition directly from conditioned culture medium and found that fragment size distributions in mutant cells mimics the hypofragmented profiles previously observed in plasma samples from p.R206C carriers. We also showed that *in vitro* treatment of hypofragmented cfDNA with recombinant wildtype DNASE1L3 could enrich for mononucleosomal fragments, with fragment end-motifs characteristic of DNASE1L3 cleavage activity. This could open avenues for DNASE1L3 as a candidate pre-treatment agent to improve the accuracy and efficiency of cfDNA sequencing-based diagnostics in hypofragmented liquid biopsies. These findings demonstrate that our isogenic cell line model provides a controlled system to study cfDNA fragmentation biology and DNASE1L3 function.

## Introduction

Cells release extracellular or cell-free DNA (cfDNA) upon cell death into the bloodstream, providing a powerful biomarker for non-invasive clinical diagnostics and disease monitoring.^1,2^ cfDNA analysis supports diverse applications in liquid biopsy testing: for instance cell-free tumor DNA (ctDNA) captures the landscape of genomic heterogeneity in tumor genomics^3^, while cell-free fetoplacental DNA (cffDNA) is used for non-invasive prenatal testing (NIPT) to monitor fetal aneuploidies.^4^ The emerging field of *cfDNA fragmentomics* studies fragmentation patterns, using metrics such as fragment size, fragment end compositions, and epigenetic markers.^5^ Such fragmentomic markers can be used to identify patterns that correlate with disease and to improve diagnostic accuracy.^1^ By studying the underlying mechanisms of cfDNA biology, fragmentomics not only provides fundamental insights in enzymatic steps involved, but may also offer insights on how liquid biopsy approaches can be improved.^6^

Aberrant cfDNA homeostasis can compromise liquid biopsy testing performance.^7,8^ Fragmentomic studies have shown that cfDNA undergoes non-random fragmentation through a cascade of deoxyribonucleases (DNases) that occurs both intracellularly and systemically.^1,9^ Plasma cfDNA fragment length distributions typically display a major peak around ~166 base pairs, aligning with the nucleosomal architecture of histone-bound DNA. This step-wise nucleosomal fragmentation is often referred to as apoptotic laddering^10^, and is vital for adequate formation and clearance of cfDNA.

DNase1-like 3 (DNASE1L3, also known as DNase γ, DNase Y, LS-DNase) is a major determinant of the fragmentomic properties of plasma cfDNA. It is primarily secreted by macrophages in the liver and circulating immune cells.^11^ As opposed to DNase1, DNase1L3 is not inhibited by actin, efficiently digests chromatin-associated DNA and is capable of causing apoptotic laddering on its own.^12^ Intracellularly, it digests chromatin during apoptosis^13^, while extracellularly, it helps to further degrade cfDNA in the bloodstream.^14,15^ A key mechanism in this context is NETosis, where neutrophils release neutrophil extracellular traps (NETs) composed of DNA and proteins to capture pathogens.^16^ DNASE1L3 is essential for degrading the DNA within these NETs, and failure to do so leads to the accumulation of antigenic cfDNA, triggering the production of autoantibodies against DNA and nucleosomal components and contributing to the development of autoimmune diseases such as systemic lupus erythematosus (SLE).^11,17,18^ Complete loss-of-function mutations in *DNASE1L3* result in pediatric-onset SLE with hypocomplementemic urticarial vasculitis syndrome (HUVS).^19^ Treatment or even prevention of autoimmunity in knock-out mice has recently been demonstrated with an injectable engineered DNase, complementing the loss-of-function.^20^

Genome-wide association studies have identified the *DNASE1L3* p.R206C (dbSNP: rs35677470) missense variant as a moderate risk factor for systemic autoimmune diseases such as SLE and systemic sclerosis (SSc). Homozygosity for this common variant (7% allele frequency in Europeans, but lower in other populations) significantly increases the likelihood of inconclusive non-invasive prenatal testing (NIPT) results due to low cfDNA recovery yield.^7^ cfDNA in such samples is hypofragmented and presents lower plasma cfDNA concentrations, compared to that of controls, as a result, too little cfDNA can be obtained from plasma to perform a sensitive NIPT. The p.R206C mutation disrupts the protein’s tertiary structure, impairing both its catalytic activity and secretion into the bloodstream.^15^ Previous studies have utilized murine models to investigate DNASE1L3’s role in fragmentomics and systemic autoimmune disease.^21–24^ However, these models are time-consuming and resource-intensive, and need to be confirmed by a human model, delaying progress. Although relatively time- and cost-efficient, cell line models with exogenous plasmid-based *DNASE1L3* overexpression^25^ do not properly reflect natural expression profiles. Such constitutive, “always on”, expression models do not accurately represent the reported intracellular “stand-by level” physiology of DNASE1L3, complicating its characterization under native apoptosis-dependent triggers. Furthermore, the transient nature of such expression models limits their use in longer-term cultures, as they require selection pressure in the form of antibiotics that can directly affect metabolism and growth.^26^ Hence, it would be preferable to study p.R206C in *DNASE1L3* in a stable isogenic context, keeping all other genetic factors identical.

Genome editing using Clustered Regularly Interspaced Short Palindromic Repeats (CRISPR) and CRISPR-associated (Cas) proteins has revolutionized genetic research, enabling targeted modification of genomic DNA. Traditional knock-in methods using Cas9, involve introducing a double-strand break (DSB) and a donor oligonucleotide containing the variant of interest. However, DSBs often lead to undesired insertions, deletions or sometimes even translocations, complicating the introduction of a single point mutation. Recent developments in high-precision CRISPR-based genome editing have enabled efficient single-base modifications by using a Cas9 nickase fused to a reverse transcriptase (RT). Dubbed prime editing, this technique employs a chimeric prime editing guide RNA (pegRNA) that combines a CRISPR RNA (crRNA) and an RNA RT template carrying the desired knock-in sequence.^27^

We here used CRISPR prime editing to develop an isogenic cell line model to study DNASE1L3 p.R206C *in vitro*. This cell line model can be used for studying fragmentomic features of cfDNA secreted into conditioned medium. Using this model, we confirm the effect of p.R206C on fragmentomic properties, and we show that adding recombinant DNase1L3 partially restores fragmentation patterns. This system provides a controlled platform to investigate DNASE1L3 function in cfDNA fragmentation and offers insights into potential strategies for mitigating the effects of cfDNA fragmentation deficiencies in clinical applications.

## Materials & Methods

### Plasmid construction & isolation

A plasmid expressing pegRNA-DNASE1L3^R206C^, a pegRNA to introduce the *DNASE1L3* c.616C>T (dbSNP: rs35677470) mutation endogenously, was derived from pU6-pegRNA-GG-Vector (Addgene #132777) through Golden Gate assembly, as described by Anzalone and colleagues^27^. We used *DNASE1L3* transcript variant 1 (accession no. NM_004944) for pegRNA design. Plasmid construction and cloning was done in *Escherichia coli* subcloning efficiency DH5α competent cells (Invitrogen™, Thermo Fisher Scientific) in solid and liquid Luria-Bertani (LB) medium, prepared from Difco™ Agar Noble (Becton, Dickinson and Company) and Difco™ LB broth, Lennox (Becton, Dickinson and Company) respectively. Constructs were heat-shock transformed at 42°C for 40 seconds, followed by immediate cool down on ice for 5 minutes and plating on selective solid media. Inoculation of selective liquid media was done from a single colony. For plasmid selection, media were complemented with 100 mg/L ampicillin. Bacterial cultures were incubated at 37°C, in a Unitron® plus incubator shaker at 200 rpm (Infors HT) or a Heraeus stationary incubator (Thermo Fisher Scientific). Plasmids were purified from liquid bacterial cultures using the PureYield™ Plasmid Midiprep System kit (Promega, Madison, USA) according to the manufacturer’s instructions. Whole Plasmid Sequencing was performed by Plasmidsaurus (Plasmidsaurus) using Oxford Nanopore Technology with custom analysis and annotation.

### Prime editing in HEK293T cells

To introduce the p.R206C mutation in *DNASE1L3*, HEK293T cells (ATCC: CRL-3216™) were reverse co-transfected with pegRNA-DNASE1L3^R206C^ (see above) and pCMV-PE2 (Addgene #132775) using Lipofectamine™ 3000 (Invitrogen™, Thermo Fisher Scientific). For each transfection, two solution were prepared: Solution 1 (125 µL Opti-MEM [Gibco®, Thermo Fisher Scientific], 1 µg pegRNA-DNASE1L3^R206C^, 1.5 µg pCMV-PE2, and 5 µL P3000 reagent [Invitrogen™]) and Solution 2 (125 µL Opti-MEM, 3.5 µL Lipofectamine 3000). Both solutions were combined to a total volume of 250 µL, incubated at ambient temperature for 15 minutes and transferred to a well in a 6-well plate. 8×10^5^ HEK293T cells were placed on top, followed by adding Dulbecco’s Modified Eagle Medium (DMEM; Gibco®) supplemented with 10% fetal bovine serum (FBS). Cells were incubated overnight at 37°C with 5% CO_2_, transferred to a 10 cm dish and subsequently cultured for five days. Genomic DNA was then extracted from approximately 8×10^4^ cells using the MightyAmp™ Genotyping Kit (Takara Bio) following the manufacturer’s instructions. PCR amplification of the targeted region was performed with Terra PCR Direct Polymerase Mix (Takara Bio) using primers KD145 and KD146 with a melting temperature (Tm) of 70°C. PCR products were treated with ExoSAP-IT™ PCR Product Cleanup Reagent (Thermo Fisher Scientific) and subsequently analyzed via Sanger sequencing using primer KD146 and the Mix2Seq Kit (Eurofins Genomics) according to the manufacturer. Sequencing results were analyzed in SnapGene® version 7.0.3 to confirm the presence of the p.R206C mutation in the pool. To generate single-cell clones, cells were seeded at a density of 0.5 cells per well in 96-well plates and cultured up to 25-50% confluency. Next, each well was split into two fractions: one fraction was used for propagation of the clone, the other was used for PCR and Sanger-based verification of the p.R206C edit as described above. Successfully edited clones were transferred to flasks for further growth.

### qPCR

Total RNA was extracted from approximately 1×10^6^ HEK293T cells using the High Pure RNA Isolation Kit (Roche), following the manufacturer’s protocol. RNA concentration and purity were determined using a NanoDrop spectrophotometer (Thermo Fisher Scientific). Complementary DNA (cDNA) was synthesized from 1 µg of total RNA using the iScript™ cDNA Synthesis Kit (Bio-Rad) according to the manufacturer’s instructions. qPCR was performed using SYBR Green I Master Mix (Roche) on a LightCycler® 480 System (Roche). Each reaction contained 1 µL of the crude cDNA mixture and primers KD155 and KD156, with amplification performed according to the manufacturer’s cycling conditions. To prepare PCR products for sequencing, reactions were treated with ExoSAP-IT™ PCR Product Cleanup Reagent (Thermo Fisher Scientific) and heat-inactivated according to the manufacturer’s protocol. A 1:1000 dilution of the purified product was subjected to Taq PCR amplification using primers KD155 & KD156 with a Tm of 50°C. The resulting products were again treated with ExoSAP-IT™ before being submitted for Sanger sequencing with the Mix2Seq Kit (Eurofins Genomics).

### Collection of cfDNA from conditioned culture medium

For the generation of conditioned medium, 2×10^6^ HEK293T cells were seeded in 10 mL of DMEM supplemented with 10% FBS, and incubated for 10 days at 37°C with 5% CO_2_. The medium was harvested by transferring it to a centrifuge tube and spinning at 286 *g* for 5 minutes to pellet cellular debris. The supernatant was filtered through a 0.45 µm filter to remove remaining particles and stored at −20°C until further use. cfDNA was isolated as described previously.^28–30^ Briefly, automated nucleic acid extraction from thawed conditioned medium was done using QIAsymphony (Qiagen, Venlo, The Netherlands). Isolates were quantified using the Qubit dsDNA BR kit (Invitrogen, Thermo Fisher Scientific) on a Qubit 4 Fluorometer (Invitrogen, Thermo Fisher Scientific), and stored at −20°C until further use.

### Digital droplet PCR

Digital droplet PCR was done using the QX200 Droplet Digital PCR System (Bio-Rad) as previously described.^7^ In short, wild type and mutant (p.R206) alleles were detected using custom designed probes with the Bio-Rad Mutation Detection assay (Bio-Rad). Probes correspond to different fluorescence channels and were designed using the complementary online design service provided by the manufacturer. Each ddPCR reaction was done with 40 ng input DNA. Partitioning, amplification and acquisition were done according to the manufacturer.

### Fragment size analysis

For plasmid cloning construction and analysis, fragment size separation on agarose gels was done using 1% UltraPure™ agarose gels (Invitrogen, Thermo Fisher Scientific) in 1x TAE, supplemented with 0.05% (v/v) EtBr. Visualization was done using a Gel Doc XR+ imager (Bio-Rad Laboratories, Inc.). For cfDNA fragmentomic size analysis, samples were analyzed on a TapeStation 4200 (Agilent Technologies Inc., Santa Clara, USA) with D5000 DNA ScreenTape (Agilent Technologies, Inc.).

### Reconstitution assay

200 ng hypofragmented cfDNA from a *DNASE1L3* p.R206C HEK293T cell line was incubated with and without 25 pmol recombinant DNASE1L3 (Cusabio® or Aviva Systems Biology) in 1x DNase1 buffer (New England Biolabs) for 4 hours at 37°C. Reaction products were isolated with AMPure XP bead-based reagent (Beckman Coulter), with a bead:sample ratio of three, and two wash steps with 80% ethanol. Elution was done in 35µL ATE buffer (Qiagen), followed by quantification with the Qubit™ dsDNA HS kit (Invitrogen, Thermo Fisher Scientific) on a Qubit 4 Fluorometer (Invitrogen, Thermo Fisher Scientific). Isolates were diluted to 2 ng/µL, prior to fragment size analysis as described above.

### Fragment end-motif analysis

Fragment-end motif analysis was done on long-read sequencing data obtained through nanopore sequencing. Nanopore NGS library preparation was done using Nanopore V14 chemistry, with the NEBNext Companion Module for Oxford Nanopore Technologies Ligation Sequencing (New England Biolabs). Sequencing was done with Nanopore MinION Mk1B (Oxford Nanopore Technologies plc., UK). DNA isolates were barcoded with the Nanopore native Barcoding kit 24 kit (SQK-NBD114.24; Oxford Nanopore Technologies, plc.) and at least one million fragments were sequenced per sample, unless there was not sufficiently enough material. Base-calling was done with Dorado v0.9.0 SUP Model 5.0 (https://github.com/nanoporetech/dorado/releases/tag/v0.5.0). End-motifs and fragment-sizes were analyzed using the cfstats package (https://github.com/jasperlinthorst/fragmentomics) and visualized using matplotlib.

## Results

We here present a pipeline that verifies *DNASE1L3* expression in HEK293T human embryonic kidney cells, introduces the p.R206C mutation through prime editing, and subsequently uses the generated cell line to characterize fragmentomic features of cfDNA derived from conditioned medium (Figure 1).

**Figure 1.**
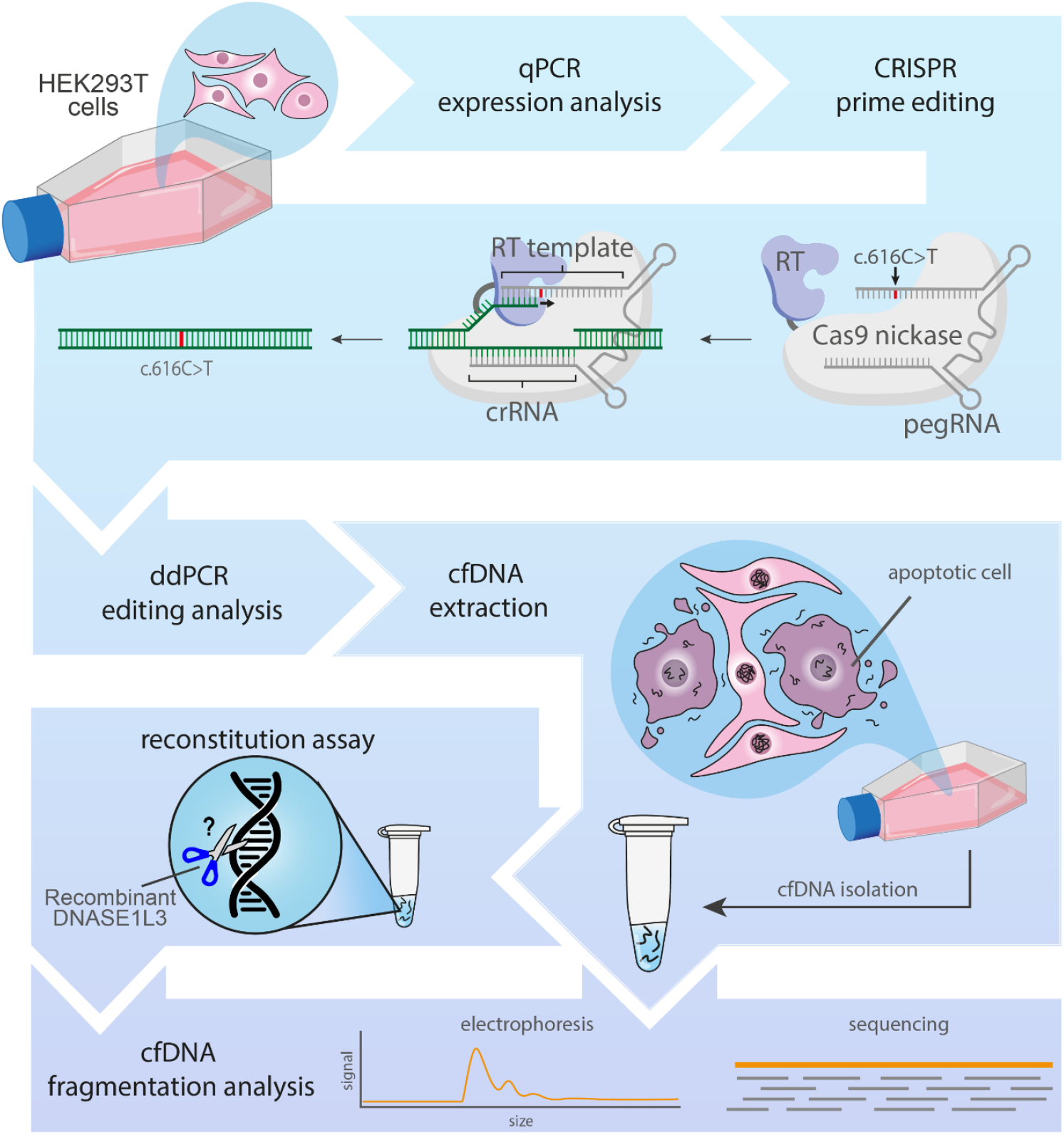
Schematic overview of the generation of a DNASE1L3 p.R206C cell line and subsequent analysis pipeline to study the fragmentation of cfDNA derived from culture medium. HEK293T cells in culture are first subjected to transcription analysis to confirm DNASE1L3 expression, followed by CRISPR prime editing. Briefly, the Cas9 nickase introduces a single-strand DNA break, while the reverse transcriptase uses a part of the pegRNA to provide a repair template in situ to incorporate the c.616C>T point mutation. DNA repair machinery attempts to repair the broken strand and mismatched base, either reverting the locus to its original state, or by mutating the other strand as well. Editing analysis through ddPCR is done to verify the correct edit. The obtained cell line model is used to extract cfDNA from conditioned medium. The fragmentation properties of this cfDNA can be studied directly through electrophoresis/sequencing, or after reconstitution treatment with recombinant DNASE1L3.

### *DNASE1L3* is expressed in HEK293T cells

Reports on expression of *DNASE1L3* in HEK293 cells (parental to HEK293T cells) diverge from 0.0 normalized RNA transcripts per million (nTPM)^31^ to positive western blot detection.^32^ Using RT-qPCR spanning exons 3 & 4 (Figure 2A), we demonstrated that, under normal culture conditions, HEK293T cells do express *DNASE1L3*, albeit at low levels. To benchmark our findings, we performed qPCR on both HEK293T and HepG2 cells, with the latter reported to express *DNASE1L3* at approximately 0.9 nTPM.^31^ Our results indicate that *DNASE1L3* expression in HEK293T cells is roughly 0.4 times that of HepG2. With Ct-values around ~35 for both cell lines (Supplementary Figure S1A), the corresponding melting curves suggest amplification of a single product (Supplementary Figure S1B). Subsequent purification and Sanger sequencing of this product further confirmed that the detected transcripts are from *DNASE1L3* (Figure 2B). Based on these results and the ease of transfection, we decided to use HEK293T cells for all subsequent experiments.

**Figure 2.**
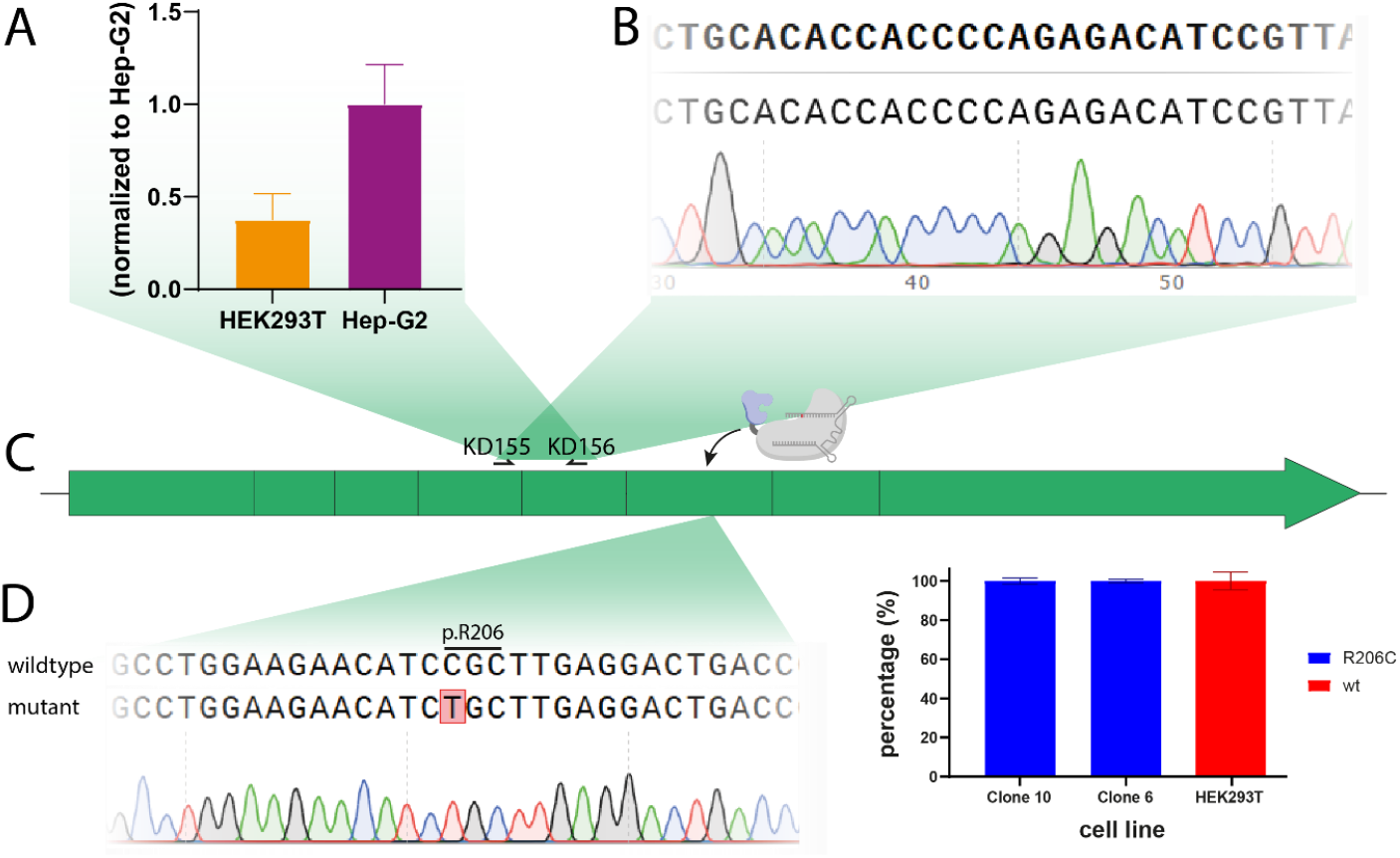
DNASE1L3 expression and mutagenesis in HEK293T cells. (A) DNASE1L3 expression in HEK293T cells, compared to Hep-G2 cells. RT-qPCR data has been normalized using TATA binding protein (TBP) as housekeeping gene. Error bars indicate standard deviation from the mean, based on triplicate reactions. Raw qPCR fluorescence and melting curve data are displayed in Supplementary Figures S1A and S1B respectively. (B) Sanger sequencing results of the obtained qPCR product map to the region that is amplified by primers KD155 & KD156 used for qPCR. (C) The Cas9 nickase prime editor was used to introduce the p.R206C mutation in exon 6. (D) Sanger sequencing of DNASE1L3 exon 6 reveals targeted editing without indels. In this case, the wild type allele [c.616C] is fully replaced by the mutant allele [c.616T], resulting in the p.R206C amino acid change in the translated product. (E) Allele distributions in a panel of prime edited HEK293T clones. Mutant (R206C) and wild type (wt) alleles were detected through ddPCR. Distributions are indicated as percentages of the total amount of droplets that detected either wild type or mutant alleles. Error bars indicate standard error of the mean of triplicate reactions. Representative raw ddPCR data is displayed in Supplementary Figure S2.

### Endogenous introduction of the p.R206C mutation in *DNASE1L3* using CRISPR/Cas9 prime editing

We transiently introduced plasmids expressing the Prime Editor complex and pegRNA into HEK293T. The pegRNA was designed to introduce the R206C variant endogenously in exon 6 of the *DNASE1L3* locus (Figure 2C). We verified correct single-base editing on single-cell clones through Sanger sequencing (Figure 2D) and ddPCR (Figure 2E).

### *DNASE1L3*^*R206C*^ causes aberrant cfDNA size derived from HEK293T cells

We have previously shown that cfDNA can be derived from cells in culture, as apoptotic cells shed fragmented DNA which can be extracted from the conditioned medium.^28,29^ The fragment size distributions of such cfDNA isolates show prominent (oligo)nucleosomal laddering patterns, mimicking plasma-derived cfDNA.^1^ Perturbations in the cfDNA fragmentation pathways, such as the R206C mutation, result in aberrant cfDNA fragment size distributions in plasma derived cfDNA.^7,9^ To see whether this could be simulated in cfDNA derived from cultured cells, we compared the size distributions of the prime edited and wild type cell lines (Figure 3).

**Figure 3.**
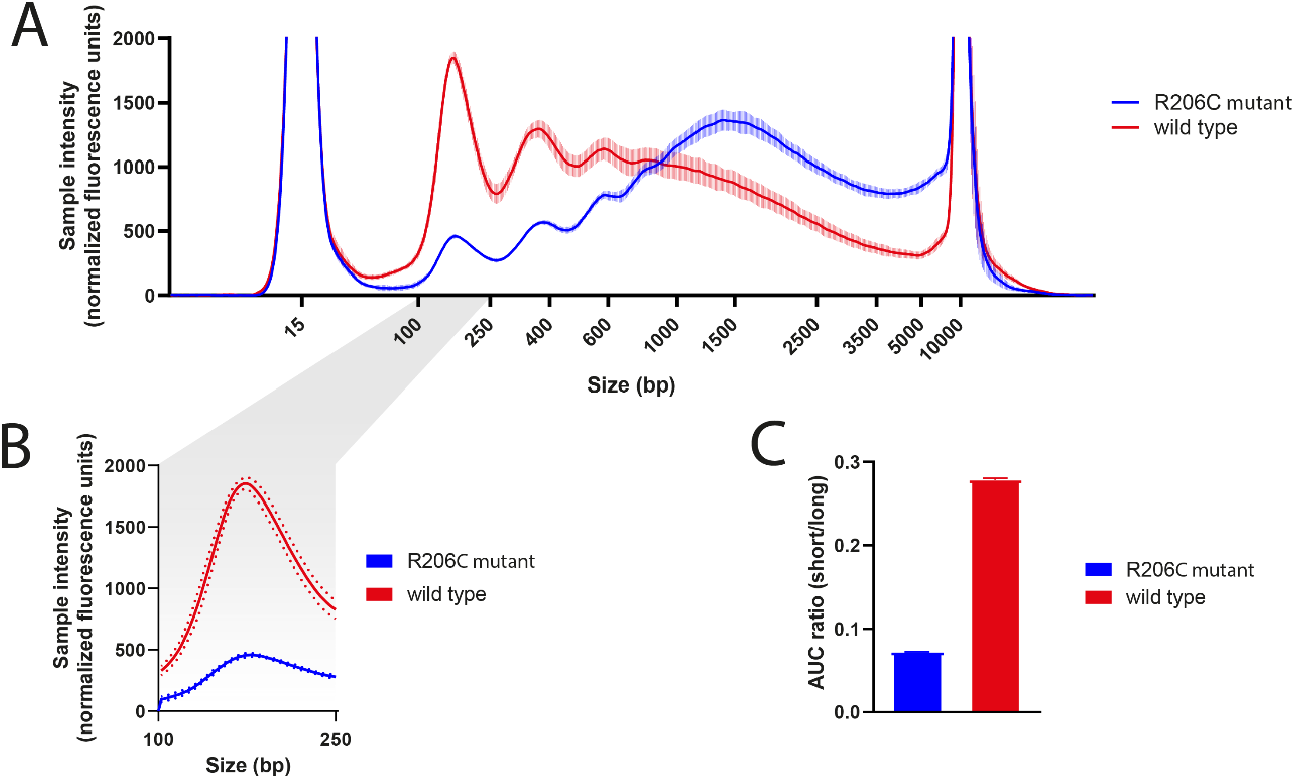
Fragment size analysis on cfDNA from conditioned medium. (A) Fragment size analysis of culture medium cfDNA from wild type and p.R206C HEK293T cells. Peaks at 15 bp and 10000 bp indicate D5000 ScreenTape lower and upper marker respectively (B) Zoom-in on mononucleosomal peaks found across the cell lines in panel [A]. (C) Ratio between the area under the mononucleosomal curve (borders 100 bp – 250 bp) and the area under the curve for fragment sizes between 250 bp – 5000 bp). Error bars and dotted lines indicate standard error of the mean of triplicate reactions.

cfDNA derived from wild-type HEK293T cells predominantly consists of mononucleosomal fragments that are between 160 and 200 bp in length (as measured by Nanopore sequencing and Tapestation), with progressively lower abundance observed for longer fragments, corresponding to the nucleosomal periodicity (Figure 3A; Figure 3B). In contrast, cfDNA from p.R206C mutant cells show a 75% drop in mononucleosomal peak height and an accumulation of fragments larger than 1000 bp. These findings suggest that the mutant DNASE1L3 produced by the prime edited cells has a reduced capacity to digest large cfDNA fragments into single nucleosomal-sized fragments. As a result, the ratio between short (100 – 250 bp) and long (250 – 5000 bp) fragments is significantly higher in wild type cells (Figure 3C).

### Functional Reconstitution of cfDNA Digestion Using Recombinant DNASE1L3

The observed hypofragmented cfDNA patterns in cell lines carrying the p.R206C variant show similarities to those reported in the plasma of individuals harboring the same mutation.^7,33^ To investigate whether the aberrant size distributions can be restored, we performed a reconstitution assay by incubating DNASE1L3 with cfDNA derived from our developed cell line model (Figure 4A). We compared two commercially available recombinant DNASE1L3 enzymes (from Cusabio® and Aviva Systems Biology), as well as DNase1, a frequently used positive control for DNA digestion. Both DNASE1L3 and DNase1 are Mg^2+^_−_ and Ca^2+^_−_dependent. After incubation at 37 °C, we isolated the DNA and performed size analysis through electrophoresis.

**Figure 4.**
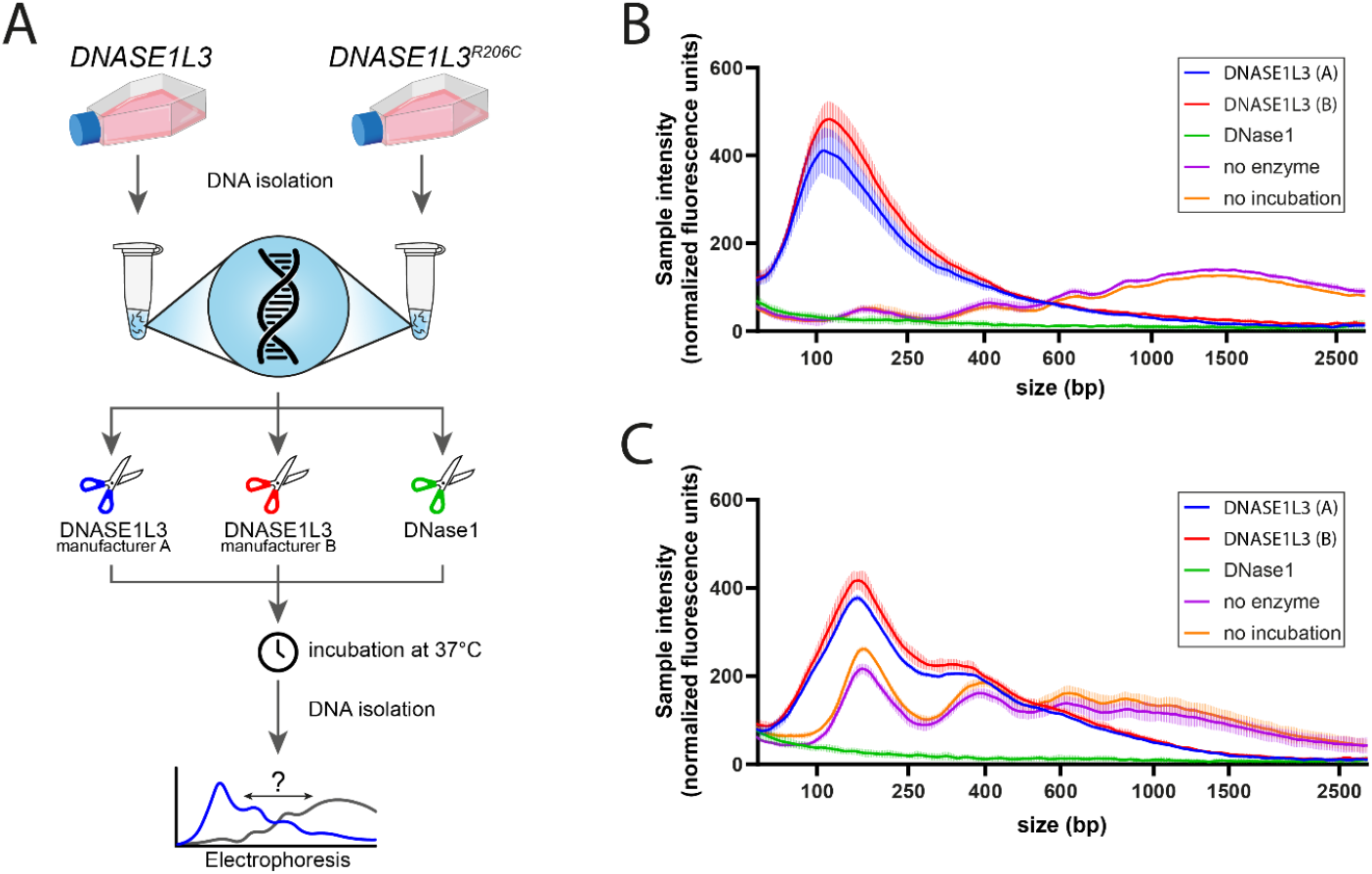
Functional reconstitution of cfDNA digestion using recombinant DNASE1L3. (A) Schematic overview of a workflow to test for artificial fragmentation of cell line-derived cfDNA using recombinant DNASE1L3 from manufacturer A (Cusabio®) and manufacturer B (Aviva Systems Biology). (B) Electropherogram for fragmentation of cfDNA from a p.R206C mutant cell line. (C) Electropherogram for fragmentation of cfDNA wild type HEK293T. “no enzyme” indicates reactions that were incubated without addition of enzyme; “no incubation” indicates control reactions that were not subjected to incubation at 37°C. Curves depict mean values of triplicate reactions, with error bars displaying the standard error of the mean.

Incubation of cfDNA from p.R206C mutant HEK293T cells with recombinant DNASE1L3 resulted, regardless of the manufacturer, in a significant shift from multinucleosomal fragment sizes to predominantly mononucleosomal fragments (Figure 4B). This demonstrates that the addition of recombinant wildtype DNASE1L3 does restore “normal” nucleosomal fragmentation. By contrast, treatment with DNase1 led to extensive cfDNA digestion into smaller fragments, consistent with its known activity to release di-, tri- and oligonucleotide cleavage products.^34^ Samples with no enzyme added, regardless of being incubated at 37 °C or not, showed no degradation, confirming cfDNA stability under these experimental conditions. To evaluate whether DNASE1L3 also affects size distributions of cfDNA from wild type cells, we ran the reconstitution assay with cfDNA from wild type HEK293T (Figure 4C). As seen with the mutant cell line, DNASE1L3 treatment also enriched the mononucleosomal population, but to a lesser extent.

### Reconstitution with recombinant DNASE1L3 enriches for mononucleosomal fragments with cytosine-rich end motifs

Murine models and GWAS on human plasma DNA screening showed that *DNASE1L3* knock-out or the p.R206C variant cause a shift not only in fragment length distribution, but also in the observed end-motifs of the fragments. In particular cytosine-rich preferences are a signature of DNASE1L3 endonuclease activity.^7,14^ To see whether the end-motifs of the samples from the *in vitro* reconstitution assay match these effects, we subjected them to long-read next-generation sequencing.

The 5’ dinucleotide end-motif distribution for different fragment lengths in cfDNA from wild type and p.R206C mutant cells were determined, each with and without recombinant DNASE1L3 treatment (Figure 5A). In accordance with the electrophoresis data (Figure 4), untreated mutant cfDNA (Figure 5A, bottom left panel) displayed a broad, right-skewed fragment size distribution, reflecting a relative accumulation of longer fragments compared to untreated wild type cfDNA (Figure 5A, top left panel). We did not observe large shifts in the usage of dinucleotide 5’ end patterns between the mutant and wildtype cell lines before treatment with recombinant DNASE1L3 (Figure 5A, top left and bottom left panels). We found that, after incubation with recombinant DNASE1L3, both cfDNA samples display very similar size and end-motif distributions, with slight enrichment for CC-ends in the mononucleosomal range (Figure 5A, top right and bottom right panels). Notably, cfDNA from wild type cell lines is also digested further to mononucleosomal size, but not smaller (Figure 5A, top right panel).

**Figure 5.**
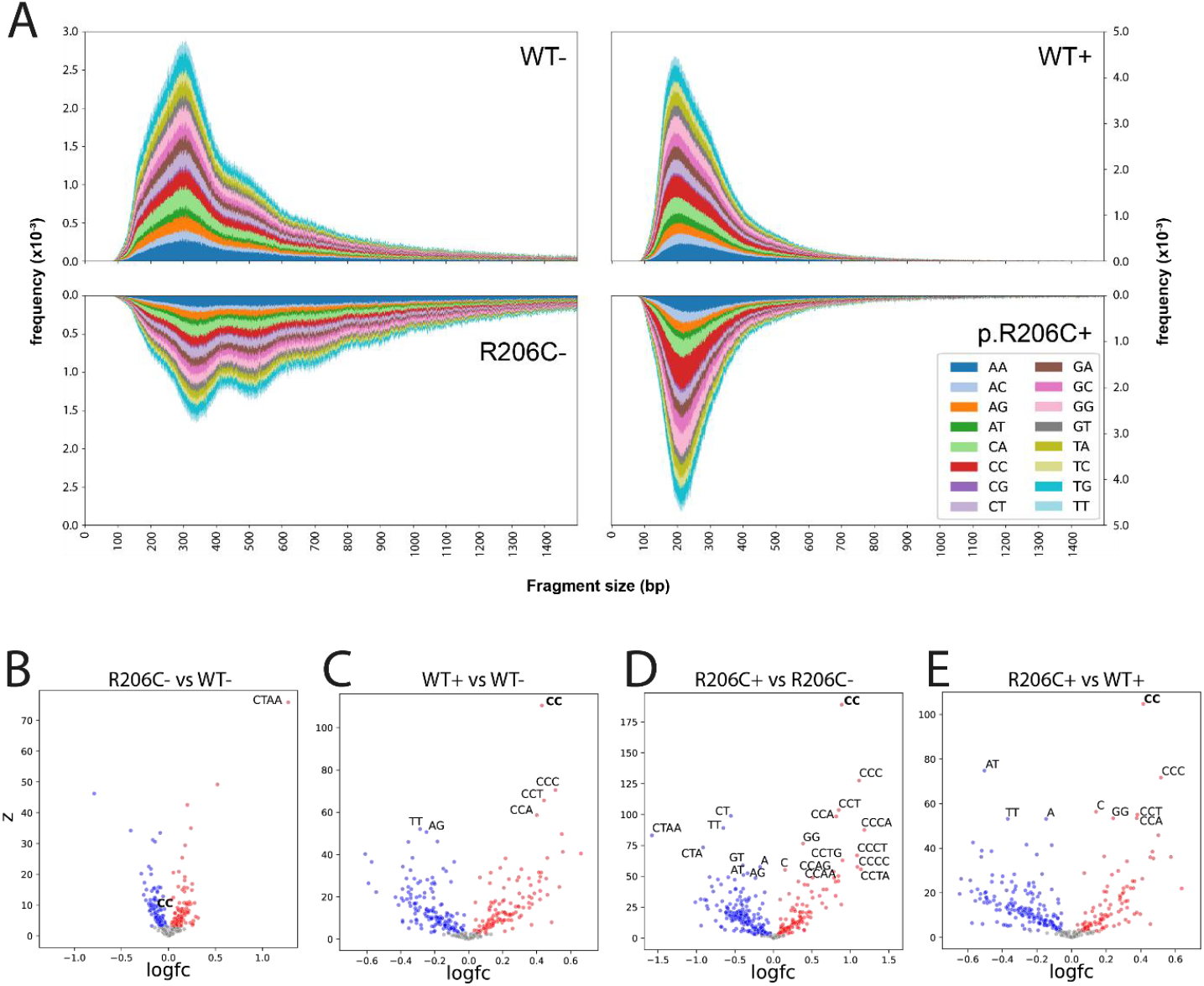
End-motif analysis on cfDNA before and after reconstitution with recombinant DNASE1L3. (A) Dinucleotide 5’ end-motif distributions in untreated wild type (WT-), untreated mutant (R206C-), recombinant DNASE1L3-treated wild type (WT+) and recombinant DNASE1L3-treated mutant (R206C+) cfDNA, for every fragment length between 60-610 bp. (B) Volcano plot displaying logarithmic fold changes in mono-, di, tri and tetranucleotide 5’ end-motifs observed in untreated mutant cfDNA, compared to untreated wild type cfDNA. Other volcano plots display the end-motif fold changes in (C) treated wild type cfDNA, compared to untreated wild type cfDNA; (D) treated mutant cfDNA, compared to untreated mutant cfDNA; (E) untreated mutant cfDNA, compared to treated wild type cfDNA. CC end-motifs and end-motifs with a Z-score above 50 are labeled. The recombinant DNASE1L3 data in this figure was obtained using Aviva Systems Biology recombinant protein. Highly similar data was obtained using Cusabio® recombinant DNASE1L3 (Supplementary Figure S3). Non-treated samples were not subjected to incubation at 37°C.

To examine sequence preferences in more detail, we compared the frequencies of end-motifs up to four nucleotides. Comparing the untreated mutant and wild type samples showed relatively heterogeneous motif usage regarding C-rich motifs (Figure 5B). In contrast, recombinant DNASE1L3-treated samples (Figure 5C-E) showed clear accumulation of cytosine-rich motifs, such as CC, CCC, CCT and CCA, in particular when treating mutant-derived cfDNA (Figure 5D). These and other C-rich sequences are among the top significantly increased end motifs in the volcano plots (e.g., upper right quadrant of Figure 5D), indicating results of specific DNASE1L3 cleavage activity. The higher magnitude of cytosine-rich motif enrichment in treated mutant cfDNA compared to treated wild type (Figure 5E) suggests the presence of more hypofragmented substrate available for cleavage, leading to a more detectable shift in motif profile upon treatment. These findings align with previous reports on cytosine-rich end-motifs of DNASE1L3 cleavage products, and underscore the suitability of our model.^35,36^

## Discussion

In this study, we developed an isogenic HEK293T cell line model to investigate the functional impact of the *DNASE1L3* p.R206C variant on cfDNA fragmentation. We first confirmed low-level *DNASE1L3* expression in the HEK293T cells we use. Next, we introduced the p.R206C variant using CRISPR prime editing, and studied the effects on cfDNA purified from the conditioned cell culture medium. cfDNA from p.R206C mutants exhibited significant hypofragmentation, with reduced mononucleosomal fragments and accumulation of larger fragments. Such hypofragmentation poses a technical challenge for next-generation sequencing (NGS), especially with short-read sequencing platforms, which are inherently biased toward short cfDNA fragments and inefficient at capturing or aligning long, high-molecular-weight DNA. We then explored the effect of adding recombinant wildtype DNASE1L3 to cfDNA extracted from conditioned cell medium, demonstrating not only an enrichment for mononucleosomal fragments, but also a shift toward cytosine-rich end motifs characteristic of DNASE1L3 cleavage. These findings underscore the impact of p.R206C on enzymatic function in human cells and suggest a potential approach to restore these defects with the goal to eventually improve diagnostic pipelines.

Cell line models can provide a controlled and reproducible system to study the specific effects of genetic variants. We here selected HEK293T cells for their ease of genetic manipulation and rapid proliferation. However, HEK293T cells are notorious for their highly unstable karyotype, sometimes even with variations between batches and passages, potentially leading to transcriptomic instability.^37^ As this may influence cfDNA fragmentation patterns, cautious interpretation of the results is needed. Additionally, as an immortalized cell line, HEK293T may not fully represent normal cellular physiology, which could impact the generalizability of the findings. After all, cfDNA fragmentation remains a process that happens not only upon apoptosis, but also in circulation, where secreted factors from different cell types play a part. Ideally, this is best evaluated in hematopoietic cell types, such as macrophages, with a natively high *DNASE1L3* expression.^11,38^ However, CRISPR prime editing of these cell types remains challenging, due to the relatively low transfection efficiency and large required constructs. Interestingly, despite the low endogenous expression of *DNASE1L3* in HEK293T cells, we observed a profound impact of the common *DNASE1L3* p.R206C variant on cfDNA fragmentation patterns, highlighting DNASE1L3’s enzymatic activity even at low expression levels. A knock-out mutant could even further support this.

Deficiency in *DNASE1L3* activity has been linked to systemic lupus erythematosus (SLE), as impaired clearance of self-DNA can trigger autoimmune responses.^11,17,18^ Recent studies also suggest a role for *DNASE1L3* in antitumor immunity, where its downregulation in tumor-infiltrating dendritic cells is associated with poor prognosis in colon cancer.^39^ While our monoculture cell line model provides insights into cfDNA fragmentomic features, studying immune-related functions of *DNASE1L3* requires more complex systems, such as co-cultures or organoids, to better mimic the microenvironment and immune interactions. Murine models can even better approach this and have been instrumental in studying *DNASE1L3* function,^21–24,39^ but remain time- & resource-intensive to generate and maintain. In contrast, our human cell line model provides a more flexible and efficient platform to investigate cfDNA biology. By showing that cfDNA fragmentation effects can be studied directly in conditioned medium cfDNA, we highlight the potential of cell line models to rapidly generate systems for studying other fragmentation factors, such as DFFB and DNase1. This adaptability positions cell line models as a complementary tool to murine systems for advancing fragmentomics research.

The majority of preceding DNASE1L3 research utilized cell lysates, conditioned media or serum to assess DNASE1L3 cleavage activity *in vitro* on plasmid DNA.^11,40,41^ By using recombinant enzyme and cfDNA isolates from human cell lines, we were able to precisely evaluate DNASE1L3 activity on its native substrate. Our findings suggest that recombinant wildtype DNASE1L3 could be added to hypofragmented samples to further digest cfDNA into mononucleosomal fragments, with an increase in cytosine-rich ends. In addition to fragment size and end motifs, our nanopore sequencing approach also enables characterization of other fragmentomic markers, such as epigenetic cfDNA signatures, which may further identify the mechanistic impact of DNASE1L3 activity.

These findings demonstrate that recombinant DNASE1L3 can effectively digest hypofragmented cfDNA into fragments of roughly mononucleosomal length. Together with the observation that the material is not further digested beyond this point, this suggests that DNASE1L3 treatment could be a valuable tool in optimizing sequencing-oriented liquid biopsy workflows. Such an approach may be particularly beneficial for improving test accuracy in cases with naturally hypofragmented cfDNA profiles, such as those observed in pregnant individuals carrying the DNASE1L3 p.R206C variant, where non-invasive prenatal testing (NIPT) has shown increased failure rates.^7^ We showed that DNASE1L3-treated cfDNA samples retained mononucleosomal-sized fragments even in the absence of endogenous DNASE1L3 deficiency, indicating that such treatment could be used broadly across samples without compromising fragment size integrity. It is remarkable, that DNASE1L3 treatment still results in fragments of approximately mononucleosomal length, even in the absence of histone proteins due to Proteinase K treatment during cfDNA isolation. Based on the obtained data in this study, we do not have an explanation for this. Furthermore, it is worth noting that the isolation procedure of cfDNA from conditioned culture medium differs substantially from plasma processing protocols, which typically involve high-speed centrifugation to remove apoptotic bodies and vesicles.^42^ The use of lower *g*-forces in combination with a 0.45 µm filter likely yields similar results, and does not undermine the observed effects of DNASE1L3 on fragmentation. However, future studies could evaluate the impact of cfDNA isolation protocols on fragmentation profiles more systematically.

The ability to normalize cfDNA profiles by using DNASE1L3 could improve both the accuracy and reproducibility of fragmentomics-based diagnostics. However, several aspects require further investigation, including optimal incubation conditions and suitable reaction buffers. For instance, DNase1 activity can be quenched in EDTA-treated samples^43^, due to chelation of divalent cations and similar effects may occur with DNASE1L3. Since EDTA is a common anticoagulant used for the collection of plasma cfDNA, this may impede the functionality of *in biopsia* treatment. Therefore, the influence of cofactors such as divalent cations (Ca^2+^ or Mg^2+^), as well as other technical considerations such as enzyme stability and potential contaminants must be carefully managed to further develop the application of recombinant DNASE1L3 in improving cfDNA-based clinical tests. Consequentially, other considerations involve testing whether treatment is most effective at the whole blood, plasma, or extracted DNA level, as well as assessing the compatibility with different sequencing technologies and potential effects on variant calling.

In parallel, other studies are exploring the therapeutic applications of administering DNASE1L3, particularly its potential to prevent autoimmunity by facilitating the clearance of extracellular DNA.^20,44^ Our successful application of CRISPR Prime Editing to introduce the p.R206C mutation also opens avenues towards corrective gene editing, for instance by reverting the mutant allele in hematopoietic stem cells of individuals with systemic autoimmune disease. In this context, bone marrow-transplants of gene edited cells could potentially restore native plasma DNASE1L3 activity, eliminating the need for *ex vivo* recombinant enzyme complementation in liquid biopsies. Altogether, our findings establish an accessible isogenic cell line model to study cfDNA fragmentation biology, and to explore DNASE1L3 as a translational tool to improve the accuracy of fragmentomics-based diagnostics.

