## Supplementary Materials for "Breaking Bad: Exploring & Complementing the Effects of the DNASE1L3 p.Arg206Cys Variant on Cell-Free DNA from an Isogenic Cell Line Model"

### Supplementary Material

#### Supplementary Figures

**Supplementary Figure S1:** DNASE1L3 qPCR fluorescence & melting curves.

**Supplementary Figure S2:** Raw ddPCR data.

**Supplementary Figure S3:** End-motif analysis on cfDNA before and after reconstitution with Cusabio® recombinant DNASE1L3.

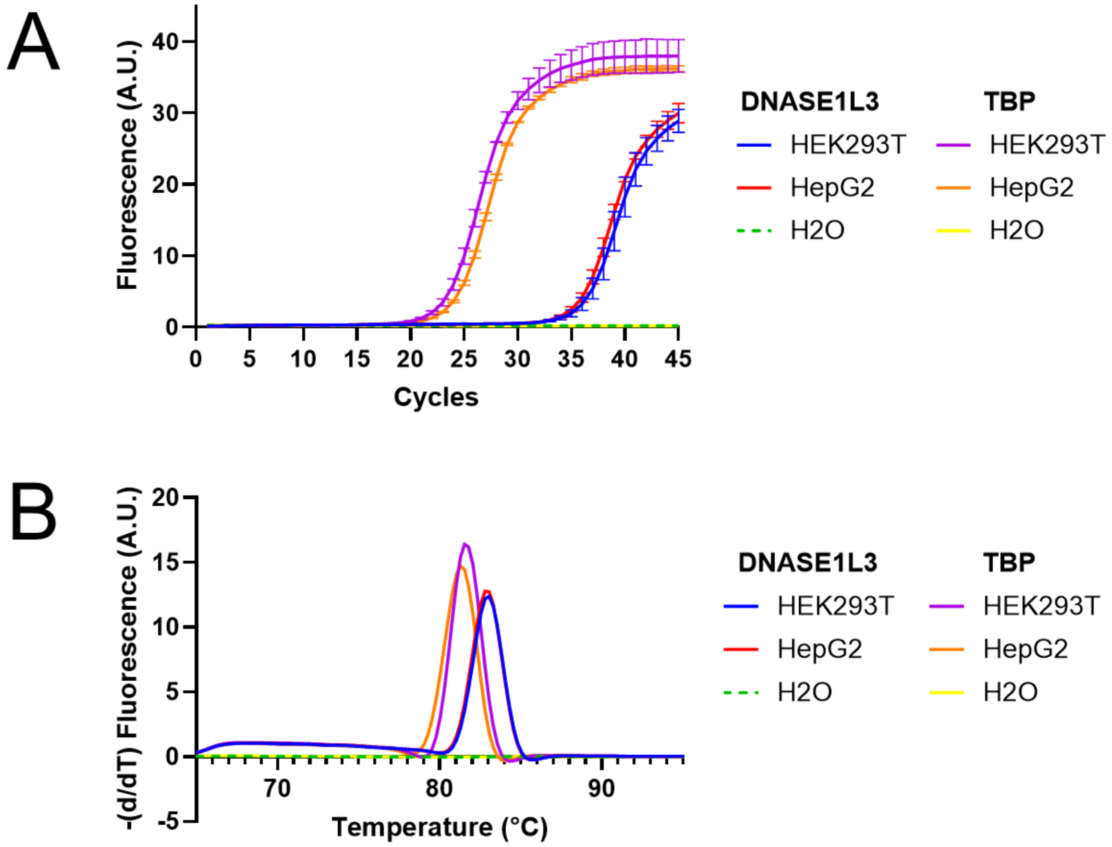


***Figure S1. DNASE1L3 qPCR fluorescence & melting curves.*** *(A) qPCR on DNASE1L3 was compared to expression of the housekeeping gene TBP in both HEK293T and HepG2 cells. Reaction with water instead of template resulted in background fluorescence only. Curves represent the mean of triplicate reactions, with error bars displaying the standard deviation. (B) Melting curves corresponding to the qPCR reactions as displayed under [A]. Curves represent the mean values of triplicate reactions.*

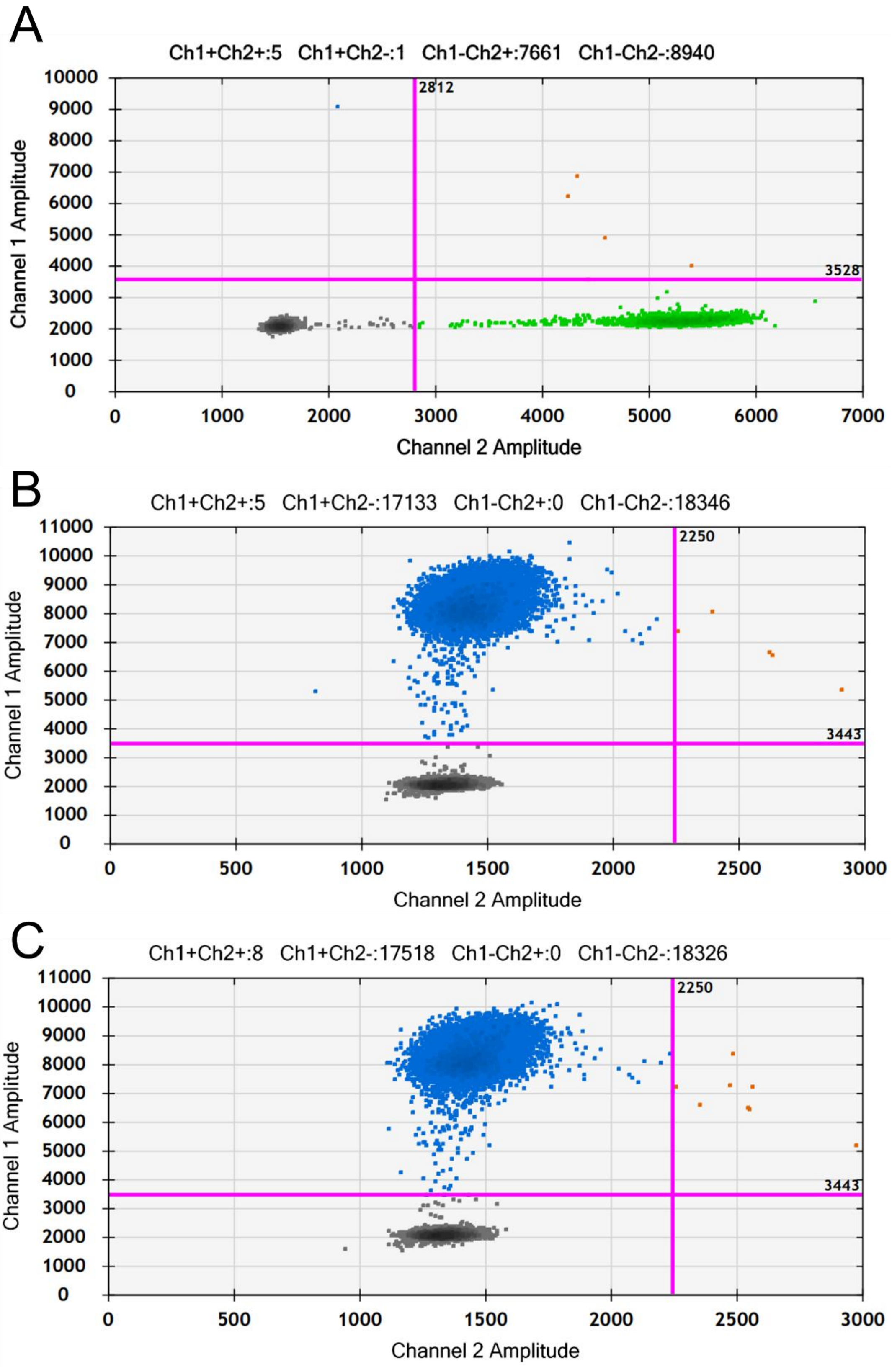


***Supplementary Figure S2. Raw ddPCR data.*** *Scatter plots from representative ddPCR replicates on cfDNA isolates from (A) wild type HEK293T, (B) mutant clone KL10 and (C) mutant clone KL6. Channel 1 represents DNASE1L3 R206C and channel 2 represents wild type DNASE1L3.*

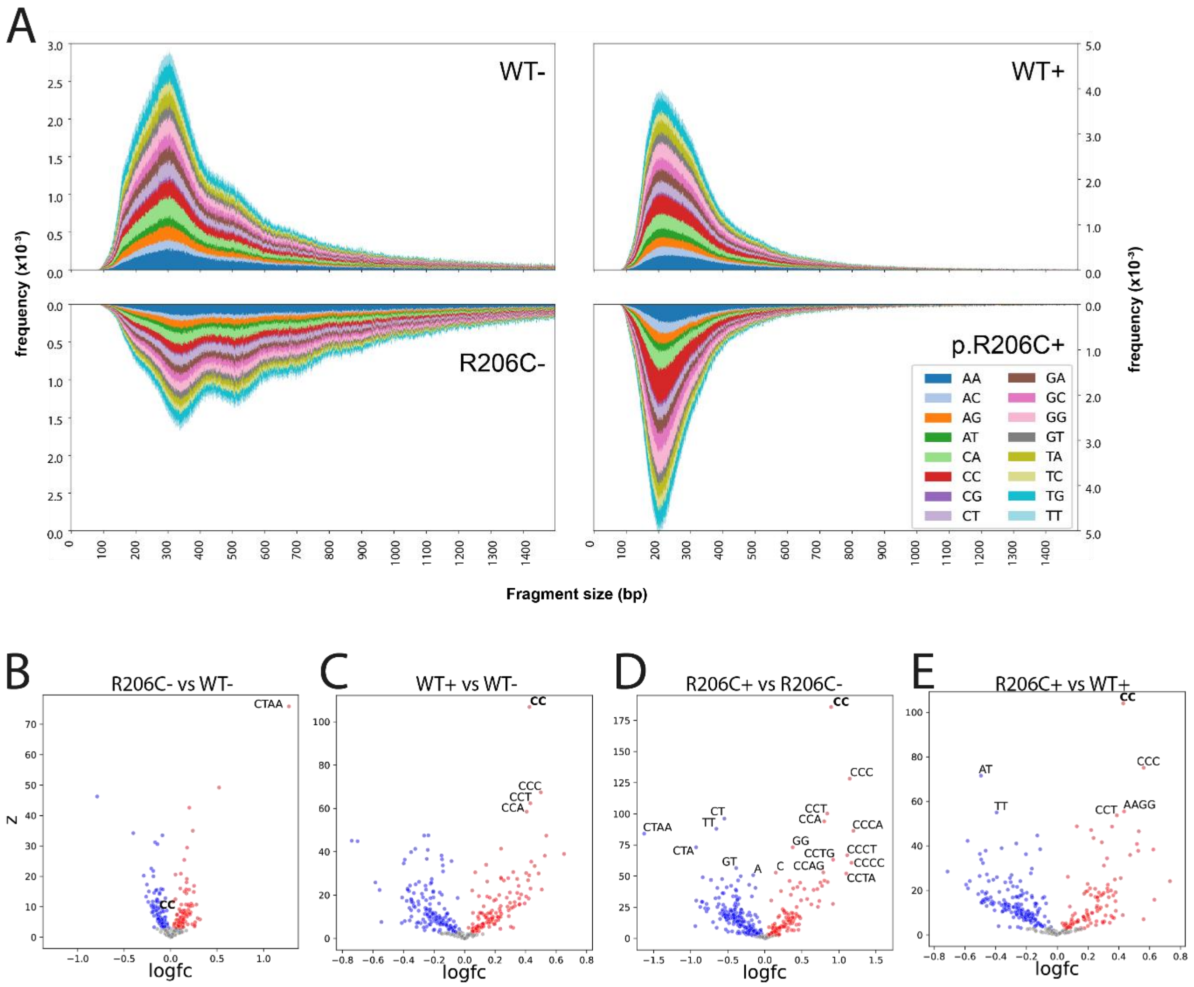


***Supplementary Figure S3. End-motif analysis on cfDNA before and after reconstitution with Cusabio® recombinant DNASE1L3.*** *(A) Dinucleotide 5' end-motif distributions in untreated wild type (WT-), untreated mutant (R206C-), recombinant DNASE1L3-treated wild type (WT+) and recombinant DNASE1L3-treated mutant (R206C+) cfDNA, for every fragment length between 60-610 bp. (B) Volcano plot displaying logarithmic fold changes in mono-, di, tri and tetranucleotide 5' end-motifs observed in untreated mutant cfDNA, compared to untreated wild type cfDNA. Other volcano plots display the end-motif fold changes in (C) treated wild type cfDNA, compared to untreated wild type cfDNA; (D) treated mutant cfDNA, compared to untreated mutant cfDNA; (E) untreated mutant cfDNA, compared to treated wild type cfDNA. CC end-motifs and end-motifs with a Z-score above 50 are labeled. The recombinant DNASE1L3 data in this figure was obtained using Cusabio® recombinant protein. Non-treated samples were not subjected to incubation at 37°C.*
